# New insights on the ventral attention network: Active suppression and involuntary recruitment during a bimodal task

**DOI:** 10.1101/782649

**Authors:** Rodolfo Solís-Vivanco, Ole Jensen, Mathilde Bonnefond

## Abstract

Reorienting attention to unexpected events is essential in daily life. fMRI studies have revealed the involvement of the ventral attention network (VAN), including the temporo-parietal junction (TPJ), in such process. In this MEG study with 34 participants (17 women) we used a bimodal (visual/auditory) attention task to determine the neuronal dynamics associated with suppression of the activity of the VAN during top-down attention and its recruitment when information from the unattended sensory modality is involuntarily integrated. We observed an anticipatory power increase of alpha/beta (12-20 Hz) oscillations in the VAN following a cue indicating the modality to attend. Stronger VAN power increases predicted better task performance, suggesting that the VAN suppression prevents shifting attention to distractors. Moreover, the TPJ was synchronized with the frontal eye field in that frequency band, suggesting that the dorsal attention network (DAN) might participate in such suppression. Furthermore, we found a 12-20 Hz power decrease, in both the VAN and DAN, when information of both sensory modalities was congruent, suggesting an involvement of these networks for attention capture. Our results show that effective multimodal attentional reorientation includes the modulation of the VAN and DAN through upper-alpha/beta oscillations. Altogether these results indicate that the suppressing role of alpha/beta oscillations might operate beyond sensory regions.

**SIGNIFICANCE STATEMENT:** Reorienting attention to unexpected events from multiple sensory sources is essential in daily life. We explored the dynamics of the ventral attention network (VAN), a set of brain regions related to attentional reorienting, when relevant information was anticipated (i.e. during top-down attention) and when unexpected congruent information from another sensory modality was presented (involuntary attentional capture). We report that activity in the alpha/beta range (12-20 Hz) within the VAN indexed both top-down and attentional capture processes. Also, the VAN was synchronized with the dorsal attention network in this frequency band, suggesting an integrated role of both networks for attentional regulation. Our results shed light on the neurophysiological mechanisms that the brain carry out for reorienting attention to relevant environmental stimuli.

## INTRODUCTION

The capacity of orienting and reorienting attention toward specific stimuli, even involuntarily, is crucial for selecting relevant information and ensuring optimal behaviour in daily life. The dorsal and ventral attention networks (DAN and VAN) have been shown to be involved in such processes as revealed by functional magnetic resonance imaging (fMRI). The DAN comprising the frontal eye fields (FEF), the superior parietal lobules (SPL), and the inferior parietal sulci (IPS), is involved in top-down attentional allocation while the VAN, encompassing the right temporoparietal junction (TPJ) and ventral frontal cortex (VFC), allows reorienting attention toward unattended relevant stimuli (see Corbetta and Shulman, 2002; Corbetta et al., 2008 for reviews; Vossel et al., 2014).

The oscillatory dynamic of the DAN and sensory areas during top-down attention tasks have also been studied using electroencephalography (EEG), magnetoencephalography (MEG) and transcranial stimulation (Worden et al., 2000; Siegel et al., 2008; e.g. Banerjee et al., 2011; Sauseng et al., 2011; Horschig et al., 2014; Marshall et al., 2015; Popov et al., 2017) while, to the best of our knowledge, very few electrophysiological studies focused on the oscillatory dynamic of the VAN during reorienting of attention (Sauseng et al., 2005; ElShafei et al., 2018; Proskovec et al., 2018). These studies have reported theta (4-8Hz) and alpha/beta (8-20Hz) decreases in the DAN and VAN during reorienting of attention to relevant stimuli, or a gamma (> 40Hz) increase in the VAN during the presentation of distracting (irrelevant) sounds to be ignored. The goal of the present study is (1) to characterize the oscillatory dynamics associated with the suppression of the activity of the VAN during top-down attention and (2) to reveal for the first time the potential additional role of this network in involuntary capture of attention across sensory modalities.

fMRI studies have revealed that during top-down attentional processes or during short-term memory involving high memory load, the activity in the TPJ is suppressed (Shulman et al., 2003; Todd et al., 2005; Shulman et al., 2007), suggesting that suppression of TPJ activity protects goal-driven behaviour from distractors. Based on the literature indicating a role of alpha oscillations in functional inhibition (Klimesch et al., 2007; Jensen and Mazaheri, 2010; Foxe and Snyder, 2011; Bonnefond and Jensen, 2012, 2013, 2015; Bonnefond et al., 2017), we predict that alpha oscillations will be high in the TPJ following the presentation of a cue directing attention to a specific modality.

Also, several studies have revealed that the VAN is involved in supramodal attention tasks (Macaluso et al., 2002; Macaluso, 2010) and it can be involuntarily activated by irrelevant stimuli coming from sensory modalities not to be attended (e.g. auditory) but spatially congruent to relevant information (e.g. visual) (Santangelo et al., 2009). In the present study, we wanted to determine whether the VAN is also recruited when information coming from an unattended sensory modality (e.g. visual) is congruent with the attended one (e.g. auditory). This process does not involve a full, voluntary reorienting of attention, as the stimulus in the attended modality remains relevant, but rather an involuntary and partial capture of attention triggered by congruency between sensory domains. Such capture should be expressed behaviourally as an enhanced performance given by congruency across modalities, instead of a cost given by attentional switch from one to the other. With a bimodal attention task, we hypothesized that the VAN would be recruited in the congruent trials of both attention conditions (visual or auditory), as reflected by a decrease of alpha oscillations in the TPJ (indicating a release from suppression, based on Solis-Vivanco et al. 2018) and/or an increase of gamma or theta oscillations (Sauseng et al., 2005; ElShafei et al., 2018; Proskovec et al., 2018). Such recruitment in congruent trials was further expected to be related to performance. In addition, we investigated the interaction between the VAN, the DAN, and sensory areas as reported in previous studies (Bonnefond and Jensen, 2012; Solis-Vivanco et al., 2018a).

## MATERIALS AND METHODS

### Subjects

We included 36 healthy subjects attending college who were recruited from Radboud University’s research participation scheme. Inclusion criteria for all participants included Dutch as their mother tongue, right-handedness according to the Edinburgh Handedness Inventory (Oldfield, 1971), normal or corrected-to-normal vision, and reported normal audition. Participants with a psychiatric or neurological diagnosis were excluded. Two participants were excluded due to excessive noise or movement artifacts during MEG recordings. The final sample consisted of 17 females and 17 males, with a mean age of 23±2.5 years. The study was conducted at the Donders Institute for Brain, Cognition and Behaviour and fulfilled the Declaration of Helsinki criteria (WMA, 2013).

### Experimental design

For this study, we re-analyzed the data from a previously reported experiment, but originally designed for both studying (1) the phase adjustment of alpha oscillations in anticipation of relevant stimuli (Solis-Vivanco et al., 2018a) and (2) the dynamics of the VAN network as described here. A cross-modal attention task was designed using MATLAB (MathWorks) custom scripts and Psychtoolbox (psychtoolbox.org). Each trial (∼5 s duration) began with a black background and a gray central fixation cross that lasted for 1 s and was projected on an acrylic screen by an EIKI LC-XL100L projector with a resolution of 1024×768 and a refresh rate of 60 Hz that lasted for 1 s (Figure 1). Subjects were asked to blink or move their eyes only during this period. Afterwards, the fixation cross turned white and 1100 ms later an electro-tactile cue (2 ms) was delivered to the left or right thumb. This cue instructed the participants to allocate attention to the visual (Attend-visual condition; 50% of trials) or auditory (Attend-auditory condition; 50% of trials) stimuli, respectively. The cue was administered with two constant current high voltage stimulators (type DS7A, Digitimer, Hertfordshire, UK; mean current=3.83 mA). After a post-cue interval of 1150 ms, visual and auditory stimuli were presented simultaneously for 200 ms, and they consisted of three syllables without meaning in Dutch. They were formed by a plosive consonant and the same vowel (‘pi’, ‘ti’, and ‘ki’). The timing of the stimuli onset and their duration was carefully controlled. The use of the same vowel (‘i’) in all stimuli further allowed us to guarantee that the length of the syllables was stable.

**Figure 1.**
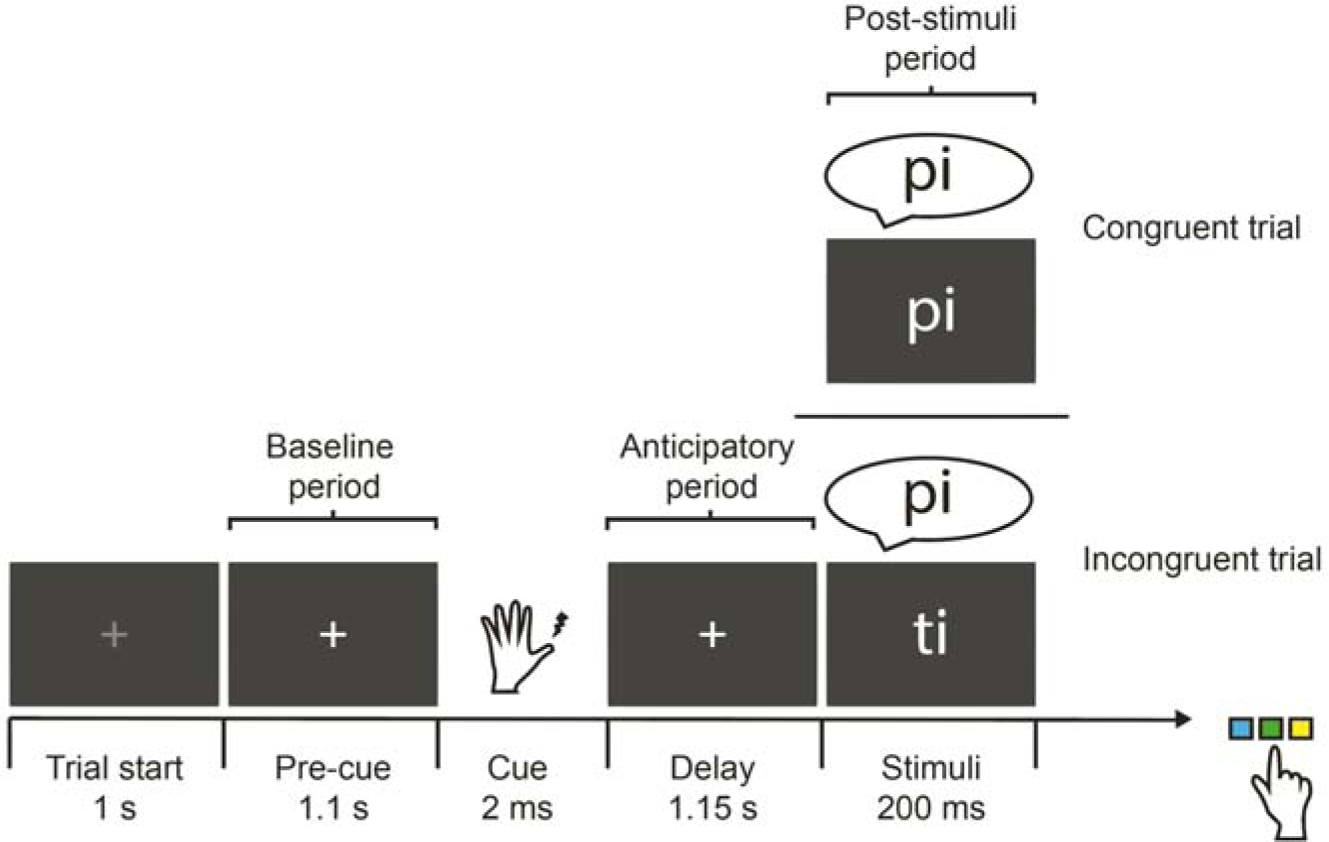
Experimental paradigm. After a lateralized somatosensory cue indicating which sensory domain to attend (visual or auditory), participants were asked to press one of three buttons according to the relevant stimulus in that domain. Baseline, anticipatory, and post-stimuli (congruency) time periods are indicated.

Each syllable was delivered with the same probability in both sensory domains. From the total number of trials (798), 75% were incongruent (different syllable between visual and auditory modality) and 25% were congruent (same syllable in both modalities). The higher number of incongruent trials was originally planned to promote anticipatory suppression of distracting information from the irrelevant sensory modality, as this was the main objective of our previous study. Nevertheless, congruent trials were included in order to explore VAN recruitment. Moreover, this proportion resembles oddball tasks used to explore TPJ activation by infrequent stimuli (Corbetta and Shulman, 2002). Visual stimuli were presented at the center of the screen in white. Auditory stimuli were digitally created using a male voice and delivered via ear-tubes adapted to MEG recordings. Each syllable was associated with either one of three buttons in a response pad. Participants were asked to respond as accurate and fast as possible to the syllable in the modality they were instructed to attend in each trial, by pressing the corresponding button using their right index, middle, or ring finger. The correspondence between the side of the cue and the modality to attend, and the assigned syllables to the buttons were counterbalanced across participants. All trials were randomly distributed across participants. Five breaks were introduced in the experiment, in which participants were informed about their performance. Reaction times (RT) and response accuracy were recorded along the experiment.

### Data acquisition

We used a whole-head magnetoencephalography (MEG) system with 275 axial gradiometers (VSM/CTF systems, Port Coquitlam, Canada) housed in a magnetically shielded room. MEG recordings were sampled at 1200 Hz with an online 300 Hz low-pass filter. The signal was down-sampled to 600 Hz afterwards for off-line analysis. No additional bandpass filtering was applied for any analysis. All participants were recorded in the supine position. Coils placed at the nasion and the left and right ear canals were used to measure participants’ head location relative to the MEG sensors during the experiment. During the recordings, an Eyelink 1000 eye tracker (SR Research, Ontario, Canada) was used to monitor eye movements and blinks.

In addition to the MEG recordings, a structural magnetic resonance image (MRI) of the participants’ brain was acquired using a 3T Siemens Trio system (Erlangen, Germany) and with a voxel size of 1 mm^3^. During the MRI acquisition, earplugs with a drop of Vitamin E in place of the coils were used for co-registration of the MRI and MEG data. Additionally, we used a FASTRAK device (Polhemus, Vermont, USA) to record the head shape of participants with 300 head points relative to the three fiducial points (nasion and the left and right ear canals).

### Procedure

The experiment was conducted over three sessions for all participants. During the first session, inclusion criteria were confirmed, general information about the study and informed consent letters were provided, and detailed instructions about the experiment were presented. Participants then performed a practice session with 150 trials inside the MEG room. During the second session, the participants’ head shape was digitized, and the actual MEG experiment was conducted. During the third session, the MRI was obtained. All data are available by request to the authors.

### Data analysis

All data analyses were done using MATLAB custom scripts and the Fieldtrip toolbox (Oostenveld et al., 2011). Epochs of the MEG recording extending 2 s before and 1 s after the onset of visual and auditory stimuli were extracted. Only epochs containing correct responses were considered for further signal analyses. From these, those containing eye blinks or saccades, muscle artifacts or superconducting quantum interference device (SQUID) jumps were rejected using an automatic routine based on mean z-scores across sensors exceeding a threshold given by the data variance within each participant. Additional visual inspection was applied to the remaining trials before demeaning and including them in further analyses. The mean number of trials included in the analysis was 559±110, with no significant differences between conditions (t=0.68, p=0.49). For the sensor-level analyses, planar gradients of the MEG field distribution were calculated (Bastiaansen and Knosche, 2000). We used a nearest neighbor method where the horizontal and vertical components of the estimated planar gradients were derived, thus approximating the signal measured by MEG systems with planar gradiometers. The planar gradients representation facilitates the interpretation of the sensor-level data, since the largest signal of the planar gradient typically is located above the source (Nolte, 2003).

Time-frequency representations (TFR) for power from 3 to 100 Hz were obtained using a fast Fourier transformation (FFT) approach with an adaptive sliding time window three cycles long (ΔT=3/f; e.g. ΔT=300 ms for 10 Hz), similarly to previous studies (Bonnefond and Jensen, 2012; Solis-Vivanco et al., 2018a). A Hanning taper (also ΔT long) was multiplied by the data prior to the FFT. For the planar gradient, the TFR of power were estimated for the horizontal and vertical components and then summed. The power for the individual trials was averaged over conditions and log-transformed.

### Source analysis

A frequency-domain beamforming approach based on adaptive spatial filtering techniques (Dynamic imaging of coherent sources; DICS) was used to estimate the power at source level in the entire brain (Gross et al., 2001). We obtained cross-spectral density matrices by applying a multitaper FFT approach (ΔT=300 ms; 1 Slepian taper resulting in 4 Hz smoothing) on data measured from the axial sensors. For each participant, a realistically shaped single-shell description of the brain was constructed, based on the individual anatomical MRIs and head shapes (Nolte, 2003). The brain volume of each participant was divided into a grid with a 1-cm resolution and normalized with respect to a template MNI brain (International Consortium for Brain Mapping, Montreal Neurological Institute, Canada) using SPM8 (http://www.fil.ion.ucl.ac.uk/spm). The lead field and the cross-spectral density were used to calculate a spatial filter for each grid point (Gross et al., 2001) and the spatial distribution of power was estimated for each condition in each participant. A common filter was used whenever two conditions were compared (based on the cross-spectral density matrices of the combined conditions). As for the sensor level analyses, the estimated power was averaged over trials and log-transformed. The power difference between conditions or time periods was calculated and averaged across participants. For the source reconstruction 33 subjects were included as the MRI of 1 subject was missing. All source data were estimated around 15 Hz according to the peak frequency effect observed in sensor level analyses (see Results section). The source estimates were plotted on a standard MNI brain found in SPM8.

In order to explore the oscillatory dynamics within regions of interest (ROI) of the VAN (see Results section), we used a linearly constrained minimum variance (LCMV) scalar beamformer spatial filter algorithm to generate maps of source activity on a 1cm grid (Van Veen et al., 1997). The beamformer source reconstruction calculates a set of weights that maps the sensor data to time-series of single trials at the source locations, allowing to reconstruct the signal at source level. In addition to TFR of power, we explored the functional connectivity across these reconstructed time series by means of TFR of coherence. In accordance to Nolte et al. (2004), we used the imaginary part of the coherence value, since it is less biased by power. All of our analyses were focused on the time period before the onset of stimuli (i.e. the anticipatory period, during which we expected a suppression of the VAN activity due to top-down attentional orientation) and the time period after (during which we explored VAN modulations due to a congruency effect between sensory modalities). A 600 ms time window before the onset of the somatosensory cue was used as baseline (Figure 1).

### Statistical Analysis

Since RT showed normal distributions (Kolmogorov-Smirnov Z for both modalities and conditions ≥0.55, p≥0.41), they were analyzed using repeated measures ANOVA (RM-ANOVA) with condition (Attend-visual and Attend-auditory) and congruency (congruent and incongruent) as within-subject factors. For all described RM-ANOVA, a Greenhouse-Geisser correction was used in case of violation of sphericity assumption and the Bonferroni test was used for post hoc comparisons.

Significant differences of power between time periods (i.e. anticipatory vs. baseline) or conditions (congruent vs. incongruent) at both sensor and source levels were assessed using a cluster-based non-parametric randomization test (Maris and Oostenveld, 2007). This test controls for the Type I error rate in situations involving multiple comparisons over sensors, frequencies and times by clustering neighboring sensors, time points and frequency points that show the same effect. For this analysis we included frequencies from 3 to 40 Hz (using 1 Hz increments) with an adaptive time window long enough to include at least 3 cycles in each frequency. We explored from −600 ms to the onset of stimuli for the anticipatory period, and from 200 to 500 ms after for the congruency effect period, based on observed effects at sensor level (see Figure 2A and Supplementary figure 1A and 1B). Sensors for which the t value of the difference between conditions exceeded an a priori threshold (p<0.05) were selected and subsequently clustered based on spatial adjacency, and the sum of the t values within a cluster was used as cluster level statistic. The cluster with the maximum sum was used as test statistic. By randomly permuting the data across the two conditions and recalculating the test statistic 1000 times, we obtained a reference distribution to evaluate the statistics significance of a given effect (Monte Carlo estimation). Additionally, for all source level analyses we also conducted a false discovery rate (FDR) correction. This correction allowed us to overcome some limitations of the cluster correction approach such as considering a set of connected smaller cluster (by chance) as one big cluster. Only clusters surviving both the cluster correction and the FDR were reported.

**Figure 2.**
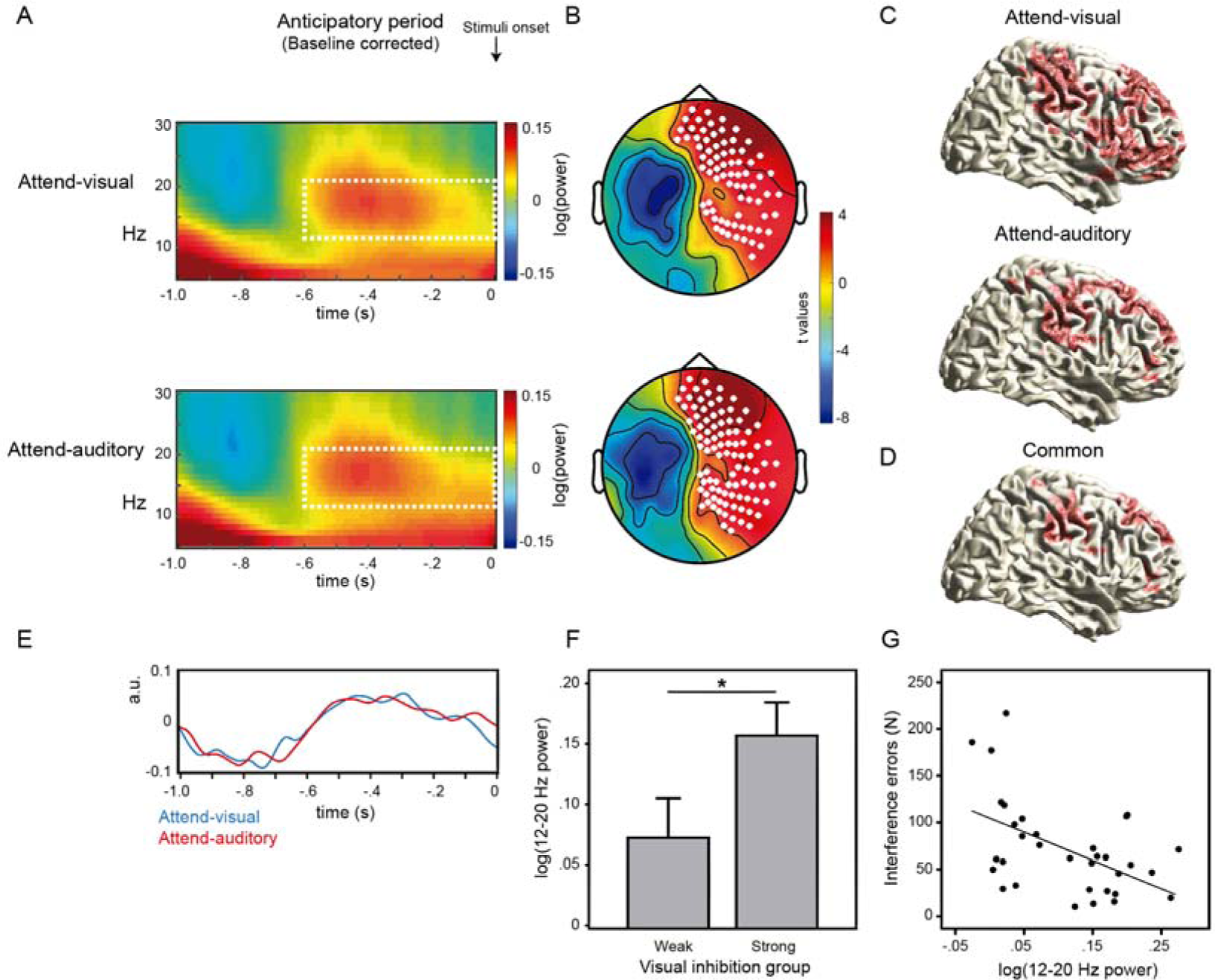
The VAN is suppressed during expectation of relevant stimuli. (A) Time-frequency representations (TFR) of power during the 1 second before the onset of bimodal (visual/auditory) stimuli for both Attend-visual and Attend-auditory conditions, compared to baseline. The dashed rectangles indicate the time-frequency window (−600 to 0 ms, 12-20 Hz) for subsequent analyses. (B) Results from cluster-based permutation tests in each condition. White dots show the sensors with significant power increases compared to baseline (p<0.05). (C) Source representations of the 12-20 Hz increase for Attend-visual (upper) and Attend-auditory (lower). Red areas indicate significant increases compared to baseline after FDR correction (p<0.01). (D) Common significant areas between both conditions. (E) Time-course of 12-20 Hz activity at source level, compared to baseline. (F) Participants with less interference errors during the Attend-auditory condition (strong visual suppressors) showed more 12-20 Hz power at source level compared to participants with more errors (weak visual suppressors) *p=0.05. (G) Higher increase of 12-20 Hz power for both conditions at source level was associated with less total number of interference errors along the task (p=0.004).

## RESULTS

We used a bimodal (visual/auditory) attentional task that included cueing for relevant stimuli (Figure 1) to quantify the neurophysiological activity associated with active suppression of the VAN activity during top-down guided attentional orientation. By controlling congruency between the sensory modalities, we also explored the recruitment of the VAN when unexpected relevant (congruent) information arising from the unattended modality was presented, and whether this improved task performance.

### Congruent stimuli enhance task performance

Behavioural results were reported before in Solís-Vivanco et al. (Solis-Vivanco et al., 2018a). Briefly, RT analysis showed that subjects were faster for the Attend-visual compared to the Attend-auditory trials (834 ± 180 vs. 919 ± 178 ms respectively, p<0.001). The RTs also showed a congruency effect, as they were faster for the congruent compared to incongruent trials for both the Attend-visual and Attend-auditory conditions (837 ± 171 vs. 947 ± 181 ms respectively p<0.001). Accuracy was better for Attend-visual compared to Attend-auditory trials (91% vs. 88%, p=0.02). Again, a congruency effect was observed, as congruent trials showed better accuracy compared to incongruent (95% vs. 83%, p<0.001).

In summary, attention was more effective for visual compared to auditory stimuli, as revealed by reduced RT and larger number of correct responses. Also, congruency between sensory modalities enhanced performance in both conditions, showing that involuntary attention capture took place.

### VAN activity is suppressed during expectation of relevant stimuli

During the anticipatory (cue-stimuli) period, we observed a power increase in a 12-20 Hz range over right scalp regions before stimuli onset, compared to baseline (−600 ms to cue onset), in both conditions (Figure 2A). The cluster-based randomization test controlling for multiple comparisons over time (baseline vs. anticipatory period), frequency (3-40 Hz), and sensors revealed that this difference was significant from 600 ms before stimuli onset in the 12-20 Hz range, regardless of condition (cluster-level statistic [CS] for Attend-visual = 3346, p = 0.014; CS for Attend-auditory = 4500, p = 0.011, Figure 2B), and remained significant when combining both conditions (CS = 4065, p = 0.018). The cluster test also revealed a significant decrease in the 12-24 Hz range on the left sensorimotor cortex (CS for Attend-visual = −108, p = 0.004; CS for Attend-auditory = −108, p = 0.001). This effect might be interpreted in terms of motor preparation, since participants always responded with the right hand (see Tzagarakis et al. (2015)), rather than an effect of the tactile cue (delivered to one thumb or the other in a counterbalanced way) so it was not further analyzed.

The right-sided power increase of 12-20 Hz was further explored at source level (15 Hz) under FDR correction (CS for Attend-visual = 915, p = 0.005; CS for Attend-auditory = 1010, p = 0.004), and revealed significance at right middle and superior frontal, temporal, and parietal regions in both conditions (Figure 2C). After identifying the common significant areas between conditions (Figure 2D) and sorting them in terms of their statistical value (t), the VAN (inferior frontal gyrus (IFG; MNI [45 30 30]) and temporo-parietal junction (TPJ, MNI [60 −52 34])), the DAN (frontal eye fields (FEF; MNI [40 - 2 50]) and superior parietal lobule (SPL; MNI [40 −50 58])), the right middle frontal gyrus (MFG, MNI [10 38 58]), and supramarginal gyrus (MNI [50 −28 36]) remained included. Also, the signal reconstruction after LCMV filter at these significant areas revealed the 12-20 Hz increase for both conditions from 600 ms before stimuli onset (Figure 2E).

In order to test whether this effect was related to task performance, we calculated the averaged power values across the grid points with the strongest effect (after FDR correction) in each condition and compared participants with low vs. high number of interference errors (i.e. responding to the unattended modality instead of the attended one) in that condition (median split). T-tests revealed that participants with less visual interference errors (strong visual suppressors) showed higher 12-20 Hz power during the anticipatory period compared to participants with more visual interference errors (weak visual suppressors) under the Attend-auditory condition (t(31)=-2.0, p=0.05, Figure 2F). This effect was not observed under the Attend-visual condition when comparing strong vs. weak auditory suppressors (t(31)=-0.56, p=0.58). We did not find a significant effect of the 12-20 Hz increase on RT for any condition. Also, we explored the association between averaged power values of both conditions and the number of total interference errors along the task. We found that stronger power (baseline corrected) during the anticipatory period was inversely correlated with performance (r=-0.49, p=0.004; Figure 2G).

When exploring the functional connectivity across relevant ROI (VAN: IFG and TPJ; DAN: FEF and SPL, selection based on power analyses at source level, see Figure 2E) during the anticipatory period, we observed an increase of coherence in the 12-20 Hz range starting from 800 ms before the onset of stimuli with respect to baseline (Figure 3A). Based on the power analyses, we selected a time-frequency window from −600 ms to stimuli onset in this frequency range and conducted a RM-ANOVA with the averaged values in each ROI and condition (factor: Attend-visual vs Attend auditory), including the baseline period as a comparison reference (factor: baseline vs anticipatory period). For this analysis, we included the visual cortex and MFG as ROI, given their significant power increase during this time window and the proposed role of MFG for connecting the DAN and the VAN (Corbetta et al., 2008). Auditory cortex was very close to TPJ, so it was excluded from the analysis due to a possible false positive result. The mean 12-20 Hz difference of power between the anticipatory and baseline periods across conditions and ROI was included as a covariate in this analysis, even when it did not show a significant main effect nor an interaction with any factor. While we did not find differences between conditions (F(1,31)=0.31, p=0.58), a significant interaction of ROI by time window (anticipatory vs. baseline, F(14,434)=5.76, p<0.0001) revealed significant increases of coherency during the anticipatory period, compared with baseline, of the TPJ with the SPL, FEF, and visual cortex (Mean difference (MD)=0.015, p=0.003; MD=0.008, p=0.02; MD=0.014, p=0.004, respectively; Figure 3A) and also a trend between TPJ and IFG (MD=0.005, p=0.09). Also, SPL and MFG showed a reduction of coherency during this period (MD=-0.009, p=0.045). Interestingly, participants with less visual interference errors during the Attend-auditory condition (strong visual suppressors) showed higher coherence between TPJ, SPL, FEF, and visual cortex compared with weak visual suppressors (F(1,31)=5.93, p=0.02; Figure 3B). In order to explore the causal relationship between DAN and VAN nodes, we calculated TFR of the phase-slope index between them for both conditions. Nevertheless, these analyses did not provide reliable results, probably due to small signal to noise ratios. No further causal analyses were carried out.

**Figure 3.**
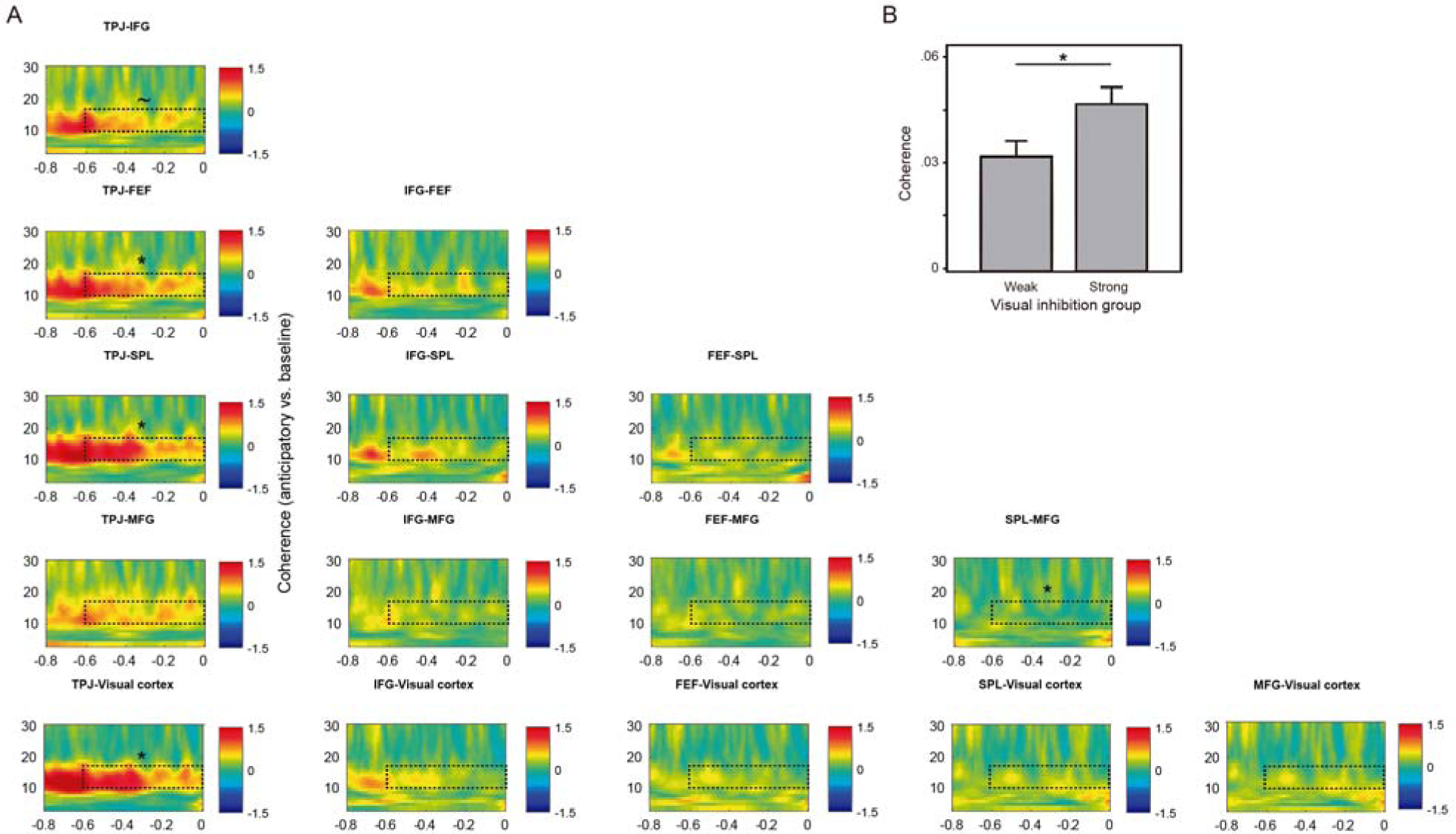
The VAN and DAN networks show increased functional connectivity during top-down attentional orienting. (A) The TPJ showed increased coherence with FEF, SPL, and visual cortex during the anticipatory period in both conditions (displayed averaged and baseline corrected), while SPL and MFG showed a reduction. The dashed rectangles indicate the time-frequency window of interest. (B) Participants with less interference errors during the Attend-auditory condition (strong visual suppressors) showed more coherence between significant nodes (TPJ, FEF, SPL, and visual cortex) compared to participants with more errors (weak visual suppressors). ∼ p<0.1; * p<0.05.

In summary, a power increase of 12-20 Hz was observed in right cortical regions before the onset of relevant stimuli, including areas from the VAN and DAN. Also, higher power increases predicted better task performance. Moreover, we observed increased functional connectivity between VAN (especially TPJ) and DAN nodes (SPL and FEF), during this period. Both increases in power and connectivity were stronger in those participants with better ability to filter out distracting visual information.

### The VAN is recruited after detection of congruency across sensory modalities

We explored whether the regions that showed 12-20Hz power increase during the anticipatory period (VAN and DAN nodes) were also modulated by attention capture, i.e. elicited by congruency between attended and unattended stimuli. We selected grid points with maximal power differences between the anticipatory period and baseline including both conditions together (though this grid points were also significant for each condition separately) and reconstructed the signal at those points during the congruency period (from stimuli onset to 600 ms afterwards) by means of an LCMV filter. These grid points included the right TPJ, IFG, FEF, and SPL. In addition, we included the visual cortex (Figure 4A). The MFG was not included in this analysis since it did not show a significant effect during congruency period (See Supplementary figure 1). TFRs of these ROIs revealed a clear decrease for congruent compared to incongruent trials in the 12-20 Hz band starting around 150 ms after stimuli onset (Figure 4B). Interestingly, this congruency effect was earlier for Attend-visual than for Attend-auditory trials. When conducting t-tests (corrected for multiple time and space points comparisons) between congruent/incongruent trials along the 0-0.5 s time window for each condition and ROI, we found significant effects in TPJ, IFG, FEF, and SPL. The TPJ and IFG were the regions that showed a congruency effect in both conditions (Figure 4C). The latency difference was further explored by comparing averaged power values of VAN ROI (TPJ and IFG) with a RM-ANOVA that included condition (Attend-visual/Attend-auditory), congruency (Congruent/Incongruent), and time window (0.1-0.3 and 0.3-0.5 s). This analysis revealed that the congruency effect was earlier for the Attend-visual condition compared to Attend-auditory (condition by congruency by time window interaction: F_(1,32)_ = 4.64, p = 0.04; Figure 4D).

**Figure 4.**
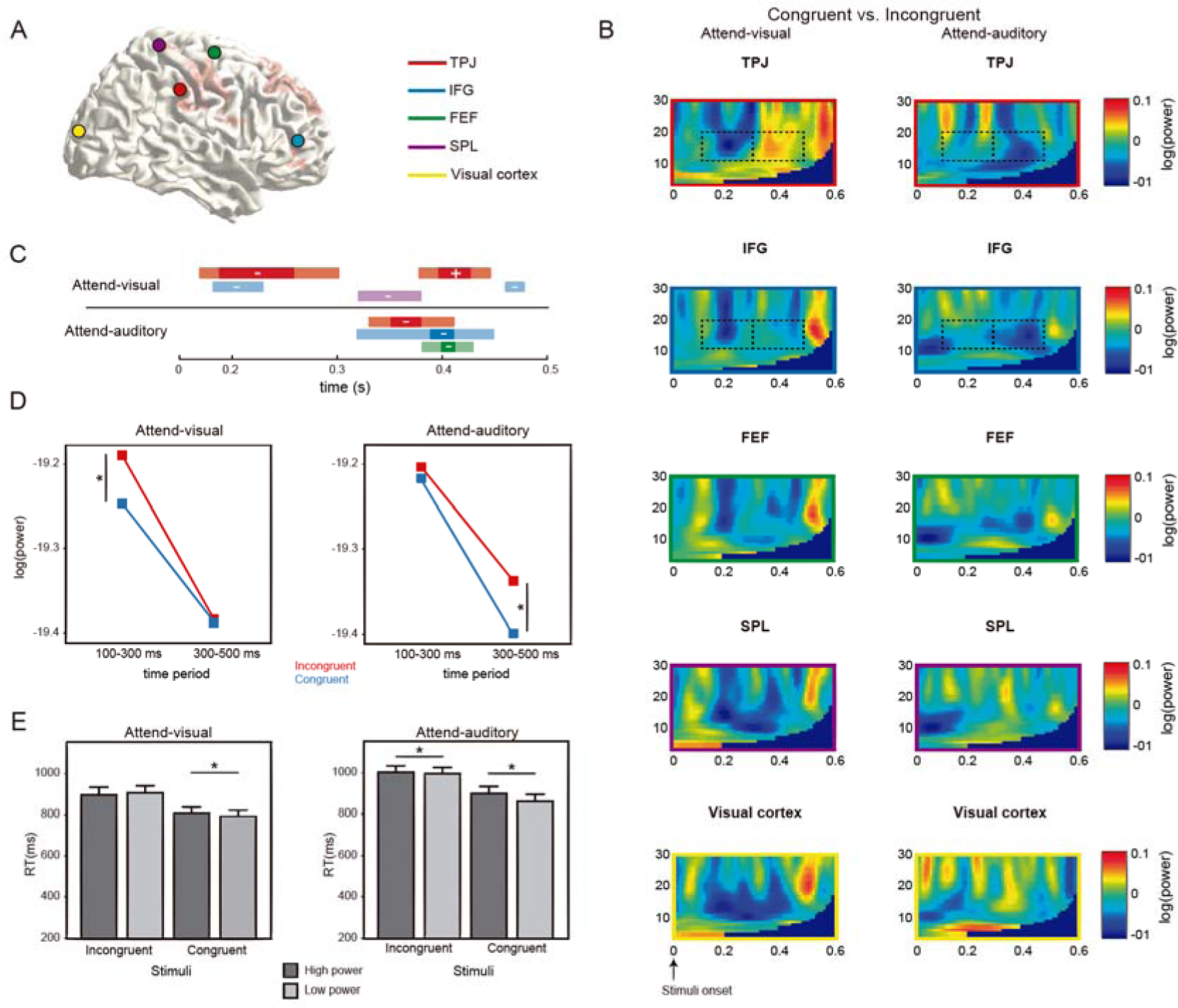
The VAN is recruited after detection of previously unattended congruent stimuli. (A) Specific ROI were selected to explore congruency effects based on the peak differences during the anticipatory period. (B) TFR of congruency effects (congruent vs. incongruent) in each ROI including right temporo-parietal junction (TPJ), inferior frontal gyrus (IFG), frontal eye field (FEF), superior parietal lobule (SPL), and visual cortex. An earlier 12-20 Hz power decrease was observed for Attend-visual vs. Attend-auditory conditions (dashed rectangles). (C) TPJ and IFG showed a significant congruency effect in both conditions during a 0-0.5 s time interval after stimuli onset. Light sections of the bars indicate significant effects (decrease (-) or increase (+)) with p<0.05. Full-color sections of the bars indicate p<0.01 (corrected for multiple comparisons in space). (D) Congruency effects in TPJ and IFG (averaged) were earlier for Attend-visual compared with Attend-auditory conditions, in accordance to time-frequency windows identified in TFR (B). (E) In both conditions, congruent trials with lower 12-20 Hz power at regions showing congruency effects were associated with reduced reaction time (RT). *p<0.05.

We further explored whether the 12-20 Hz power decrease was associated with task performance (Figure 4E). Our hypothesis was that trials with low alpha/beta power (i.e. higher activation of the VAN) would allow an enhanced congruency effect and consequently faster performances, compared with trials with high power (lower VAN activation). To this end, we classified the trials in each participant as showing low or high power in each condition and congruency variant (based on a median split) from the grid points that showed a congruency effect at source level. Then we compared the RT among power, condition and congruency factors, though ignoring the congruency main effect. A RM-ANOVA revealed that congruent trials with low power showed faster RTs compared with those with high power (Power by congruency effect: F_(1,32)_ = 6.86, p = 0.013; MD for congruent trials = −24.0, p = 0.01; MD for incongruent = −0.42, p = 0.96). Also, an interaction of condition by power was found (F_(1,32)_ = 7.03, p = 0.012). Post hoc comparisons revealed that trials in general with low power reduced RT within the Attend-auditory condition (MD for Attend-auditory = −22.1, p = 0.01; MD for Attend-visual = −2.34, p = 0.74; Figure 4E).

When assessing the power congruency effect both at sensor and source levels for the whole brain, the 12-20 Hz decrease was further confirmed at similar right sensors as for the anticipatory period (Supplementary figure 1A). The cluster-based randomization test in a 200-500 ms time window showed that this effect was especially prominent for the auditory condition at sensor level (Attend visual: CS = −16, p = 0.05; Attend auditory: CS = −41, p = 0.013; Supplementary figure 1B). When exploring this effect at source level (LCMV filter and power averaged values at 15 Hz), both conditions revealed this effect on right superior and inferior frontal, temporal and parietal areas, for both conditions (300-400 ms; Attend-visual: CS=-346, p=0.037; Attend-auditory: CS=-316, p=0.048) and with a similar pattern when considering both conditions together (Supplementary figure 1C). After an FDR correction, nodes from the VAN (TPJ and IFG) remained included. At both sensor and source levels, the topographic profiles of the right sided 12-20 Hz modulation during the anticipatory (compared to baseline) and post-stimuli (congruent vs. incongruent) periods were notably similar (Supplementary figure 1D). In order to discard that the congruency effect rather reflected differences related to the overrepresentation of incongruent vs. congruent stimuli (75 vs. 25%), we explored whether congruent trials evoked the P300 event-related field (P300m) at parietal regions due to an oddball effect (Polich, 2007). Nevertheless, this ERF was not observed for any modality (data not shown).

When exploring the functional connectivity across relevant ROI (TPJ, IFG, FEF, SPL, and visual cortex) during the congruency period, we found a significant increase of coherence for congruent vs. incongruent stimuli in the alpha-beta range (8-15 Hz), this time starting around 50 ms after stimuli onset (F(1,31)=12.47, p=0.001; Figure 5), regardless of condition. A significant interaction of congruency by pair (F(9,279)=2.56, p=0.04) revealed that this difference was particularly significant between TPJ and the rest of the ROIs (IFG: MD=0.012, p=0.01; FEF: MD=0.01, p=0.003; SPL: MD=0.01, p=0.002; visual cortex: MD=0.01, p=0.01). The mean 8-15 Hz difference of power between congruent and incongruent stimuli across conditions and ROI was included as a covariate in this analysis, even when it did not show a significant main effect nor an interaction with any factor. No associations were found between coherence in these areas and task performance.

**Figure 5.**
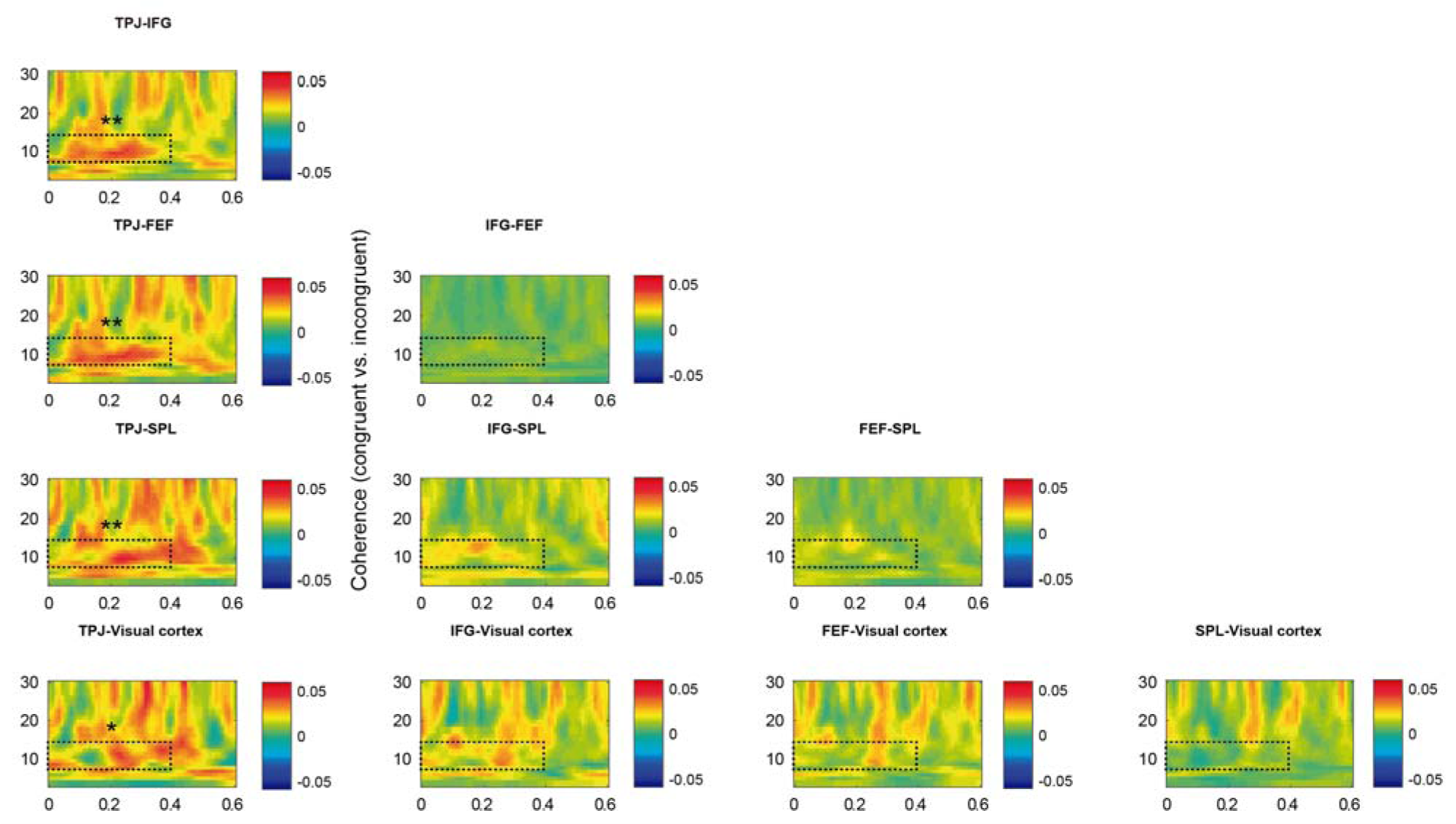
The VAN network shows increased functional connectivity after detection of previously unattended congruent stimuli. The TPJ showed increased coherence with FEF, SPL, and visual cortex after congruent stimuli onset compared with incongruent ones in both conditions (displayed averaged and baseline corrected). The dashed rectangles indicate the time-frequency window of interest. * p<0.05, ** p<0.01.

In accordance to a recent report showing an increase of gamma oscillations (>40 Hz) within the VAN after the involuntary detection of auditory distracting stimuli (ElShafei et al., 2018), we explored whether there was a congruency effect within the 50-80 Hz range after stimuli onset in the previously selected ROI. Nevertheless, no congruency effect was observed at this frequency range (Supplementary figure 2).

In summary, we found a decrease, earlier in the visual condition, in the same frequency as the anticipatory period in the VAN (TPJ and IFG), but also in the DAN (FEF and SPL), in congruent compared to incongruent trials. Such decrease predicted performance speed. In addition, increased temporal connectivity between VAN and DAN nodes was observed for congruent trials. These results suggest an involvement of the two networks reflecting effective attentional reorientation after unexpected detection of relevant information in previously unattended sensory modalities.

## DISCUSSION

In the present study, we aimed to determine (1) the oscillatory profile of the suppression of the VAN during top-down oriented attention processes and (2) whether this network was recruited when the information presented in an unattended sensory modality was congruent with the information presented in the attended modality, a special case of attention capture, in order to improve task performance. We found a power increase of alpha/beta (12-20 Hz) oscillations in the VAN during attentional orientation. This increase was associated with better task performance, suggesting that the VAN suppression might prevent shifting attention to distractors. Moreover, the TPJ was synchronized with the FEF in the same frequency band, suggesting that the dorsal attention network (DAN) might participate in the suppression of the VAN. Also, we found a 12-20 Hz power decrease, in both the VAN and DAN, when information from one sensory modality was congruent with the other, suggesting an involvement of both networks for involuntary capture of attention.

First, we observed an increase of oscillations in the 12-20Hz frequency band in VAN and DAN nodes following the presentation of the cue indicating the modality to attend, i.e. during top-down attention. Such increase was correlated with better behavioural performances, as indexed by a decrease in the number of interference errors in incongruent trials, over participants. These results suggest that the VAN is suppressed during top-down attention processes, as suggested by fMRI papers that investigated this question (Shulman et al., 2003; Todd et al., 2005; Shulman et al., 2007) and is in line with the hypothesis that such suppression would reflect a mechanism allowing to protect goal-driven behaviour from distractors. We show here that such suppression is expressed in a broad frequency band related to high alpha/low beta oscillations, i.e. in a higher frequency band compared to the frequencies observed in sensory networks. Furthermore, we observed a stronger coherence between the nodes of the VAN network as well as between FEF (part of the DAN) and TPJ. Interestingly, participants with stronger ability to suppress visual distractors showed higher connectivity across these nodes. This latest result could provide evidence in favour of the idea that the suppression of the VAN is driven by the DAN (Shulman et al., 2003). Nevertheless, since we did not find a clear direction of such connectivity between VAN and DAN (possibly due to a reduced signal-to-noise ratio), this hypothesis should be taken with caution. We also observed high coherence with sensory areas (i.e. between the visual cortex and the TPJ) during this delay period, possibly reflecting that visual information is still shared with this network in order to detect relevant information. An alternative explanation might be that as strong visual suppressors showed higher coherence including TPJ-visual cortex, this temporal communication indicates stronger suppression over sensory cortex driven by the VAN. On the other hand, we found a decrease in temporal connectivity between MFG and DAN (FEF) during top-down attention. This was unexpected, since the right posterior MFG has been discussed as another candidate region for linking the dorsal with the ventral system (Corbetta et al., 2008). Further research is needed to confirm and understand our result.

We further observed that the VAN was recruited, as indexed by a power decrease in the 12-20 Hz band, when a congruent stimulus was presented in the unattended sensory domain. Congruency effect is a special case of attention capture, as relevant information is still present in the attended dimension. Furthermore, this alpha/beta decrease predicted response speed in both conditions and was observed earlier in the Attend-visual condition, i.e. when the unattended stimulus was presented in the auditory domain, compared with the Attend-auditory condition. The processing of auditory stimuli has been shown to be faster than processing of visual ones, which could explain the earlier activation of the VAN in the Attend-visual condition (Pain and Hibbs, 2007). However, it should be noted that reaction times were faster in the Attend-visual condition, congruency effects were stronger for the Attend-auditory condition at sensor level, and VAN suppression was better in the strong visual suppressors during the anticipatory period. These results may reflect the sensory dominance of the visual domain, and hence the need for more effective modulation (suppression/recruitment) of the VAN for this type of information.

Interestingly, the DAN was also more activated during congruent than incongruent trials after an early decrease in IFG and TPJ, although only FEF reached the significant level after multiple comparison corrections in the Attend-auditory condition. In line with this, previous fMRI work has also reported higher FEF activity during reorienting of attention (e.g. Vossel et al., 2006; Corbetta and Shulman, 2011). Although the timing of activation of the different networks would need to be further investigated, it seems that the VAN was further activated by congruency earlier than the DAN, which ultimately might guide attention toward the unattended sensory domain. In line with this, Proskovec et al. (2018), reported a late synchrony in the alpha band in the DAN, possibly associated with reorienting attention, although they also reported an early activation of FEF. Interestingly, in the visual condition, we observed an increase of the oscillations in the 12-20 Hz band in the TPJ following the decrease, possibly reflecting a suppression of this node following its recruitment. The time window we could analyze did not allow to determine whether a similar increase was later observed in the Attend-auditory condition.

In addition, we observed a coherence increase between TPJ and IFG as well as between FEF and TPJ for congruent stimuli, although it was significant only in the Attend-visual condition. This alpha/beta synchrony could reveal the mechanism allowing the interaction within and between these networks during attention capture (Vossel et al., 2014), though we did not find a direct association with task performance. It should be noted that our selection of ROI for congruency comparisons was based on significant areas detected during the anticipatory period. Thus, other relevant regions involved in task performance and potentially synchronized with VAN and DAN during congruency detection may not have been considered. Future studies might explore connectivity patterns time-locked to responses after attention capture in order to elucidate this possibility.

We did not find any significant difference between congruent and incongruent trials in other frequency bands, neither in the theta band as reported by Proskovec et al. (2018) nor in the gamma band as reported by ElShafei et al. (2018). These discrepancies could be related to the paradigms across studies. Proskovec et al. used a Posner cueing task in which invalid trials (target on the uncued side) appeared 50% of the trials and could be very quickly detected. Possibly the theta increase they observed was more related to a bottom-up, locked to the stimulus processing (see e.g. van Kerkoerle et al. (2014) showing that theta oscillations might be related to feedforward activity). In our study, the mechanism of attention capture might be more complex as it requires the detection of congruency (i.e. incorporating relevance in the unattended domain while holding it in the attended one) and as mentioned above, this is a special case of attention reorienting. Although we observed a clear gamma increase in all the nodes (VAN, DAN and sensory cortex) compared to baseline, we did not observe gamma power difference between congruent and incongruent trials. Again, a difference in terms of paradigm might explain the discrepancy with ElShafei et al. study. In their paradigm, the auditory stimuli inducing a change in the VAN were not relevant to the task, unlike to the present study. It might be therefore interesting to study the role of relevance versus saliency for VAN activation in the different frequency bands. For instance, it has been reported that irrelevant, novel auditory stimuli generate a reduction of power in the alpha/beta band at parietal regions (Solis-Vivanco et al., 2018b). Nevertheless, whether the source of this decrease includes the VAN (and DAN) remains to be explored. Also, future studies might explore the role of disengagement (i.e. full switch from attended to unattended domain) over DAN and VAN activation, which was explored in the Posner task used by Proskovec et al. (2018), but not necessarily present in our study.

Among the limitations of our study, we were not able to clearly identify the auditory cortex, since it was too close to the TPJ and did not exhibit any specific modulation in this analysis nor in the ones we performed for our previous paper. Nevertheless, a recent fMRI study by Rossi et al. (2014) showed increased connectivity between auditory cortex and different nodes of the DAN and VAN during cued voluntary and novelty-driven auditory orienting, respectively. In addition, our sample included only young adults. Future research might explore the VAN and DAN modulation during attentional reorientation along development, including children and older adults. Also, how these networks can be compromised in neurologic and psychiatric disorders with attention impairment remains to be explored.

In conclusion, our results show that effective attentional reorientation, regardless of sensory modality, includes the modulation and cooperation between ventral and dorsal attention networks through upper-alpha/beta oscillations.

## ACKNOWLEDGMENTS

This work was supported by Consejo Nacional de Ciencia y Tecnología (CONACYT, grant number 261987) to R. Solís-Vivanco and by the James S. McDonnell Foundation Understanding Human Cognition Collaborative Award (grant number 220020448), a Wellcome Trust Investigator Award in Science (grant number 207550), and the Royal Society Wolfson Research Merit Award to O. Jensen. We thank Rocio Silva-Zunino, Jessica Askamp, and Paul Gaalman for their technical assistance.

## SUPPLEMENTARY CAPTIONS TO FIGURES

Supplementary figure 1. Detection of relevant information in unattended sensory modalities reduces upper alpha/low beta oscillations (A) A 12-20 Hz increase around 200 ms was observed after stimuli onset during congruent trials compared to incongruent trials. (B) Significant decreases for both conditions were observed at frontal, temporal, and right sensors. + p=0.05, *p<0.05. (C) After an FDR correction at source level, nodes from the VAN (TPJ and IFG) were observed to show power increases for congruent trials when considering both conditions together. Red areas indicate significant increases compared to congruent trial (p<0.05). (D) The topographic profiles of the right sided 12-20 Hz increase during the anticipatory (compared to baseline) and decrease during the post-stimuli (congruent vs. incongruent) periods were notably similar. Red areas indicate common significant regions between both time periods.

Supplementary figure 2. Congruency effects (congruent vs. incongruent) were not observed in the gamma range (40-80 Hz) at any ROI or condition.

## REFERENCES

1. Banerjee S, Snyder AC, Molholm S, Foxe JJ (2011) Oscillatory alpha-band mechanisms and the deployment of spatial attention to anticipated auditory and visual target locations: supramodal or sensory-specific control mechanisms? The Journal of neuroscience: the official journal of the Society for Neuroscience 31:9923–9932.

2. Bastiaansen MC, Knosche TR (2000) Tangential derivative mapping of axial MEG applied to event-related desynchronization research. Clinical neurophysiology: official journal of the International Federation of Clinical Neurophysiology 111:1300–1305.

3. Bonnefond M, Jensen O (2012) Alpha oscillations serve to protect working memory maintenance against anticipated distracters. Curr Biol 22:1969–1974.

4. Bonnefond M, Jensen O (2013) The role of gamma and alpha oscillations for blocking out distraction. Commun Integr Biol 6:e22702.

5. Bonnefond M, Jensen O (2015) Gamma activity coupled to alpha phase as a mechanism for top-down controlled gating. PLoS One 10:e0128667.

6. Bonnefond M, Kastner S, Jensen O (2017) Communication between Brain Areas Based on Nested Oscillations. eNeuro 4.

7. Corbetta M, Shulman GL (2002) Control of goal-directed and stimulus-driven attention in the brain. Nat Rev Neurosci 3:201–215.

8. Corbetta M, Shulman GL (2011) Spatial neglect and attention networks. Annu Rev Neurosci 34:569–599.

9. Corbetta M, Patel G, Shulman GL (2008) The reorienting system of the human brain: from environment to theory of mind. Neuron 58:306–324.

10. ElShafei HA, Fornoni L, Masson R, Bertrand O, Bidet-Caulet A (2018) What is in Your Gamma? Activation of The Ventral Fronto-Parietal Attentional Network in Response to Distracting Sounds. bioRxiv:469783.

11. Foxe JJ, Snyder AC (2011) The role of alpha-band brain oscillations as a sensory suppression mechanism during selective attention. Frontiers in psychology 2:154.

12. Gross J, Kujala J, Hamalainen M, Timmermann L, Schnitzler A, Salmelin R (2001) Dynamic imaging of coherent sources: Studying neural interactions in the human brain. Proc Natl Acad Sci U S A 98:694–699.

13. Horschig JM, Jensen O, van Schouwenburg MR, Cools R, Bonnefond M (2014) Alpha activity reflects individual abilities to adapt to the environment. NeuroImage 89:235–243.

14. Jensen O, Mazaheri A (2010) Shaping functional architecture by oscillatory alpha activity: gating by inhibition. Front Hum Neurosci 4:186.

15. Klimesch W, Sauseng P, Hanslmayr S (2007) EEG alpha oscillations: the inhibition-timing hypothesis. Brain Res Rev 53:63–88.

16. Macaluso E (2010) Orienting of spatial attention and the interplay between the senses. Cortex 46:282–297.

17. Macaluso E, Frith CD, Driver J (2002) Supramodal effects of covert spatial orienting triggered by visual or tactile events. J Cogn Neurosci 14:389–401.

18. Maris E, Oostenveld R (2007) Nonparametric statistical testing of EEG- and MEG-data. Journal of neuroscience methods 164:177–190.

19. Marshall TR, O’Shea J, Jensen O, Bergmann TO (2015) Frontal eye fields control attentional modulation of alpha and gamma oscillations in contralateral occipitoparietal cortex. The Journal of neuroscience: the official journal of the Society for Neuroscience 35:1638–1647.

20. Nolte G (2003) The magnetic lead field theorem in the quasi-static approximation and its use for magnetoencephalography forward calculation in realistic volume conductors. Physics in medicine and biology 48:3637–3652.

21. Nolte G, Bai O, Wheaton L, Mari Z, Vorbach S, Hallett M (2004) Identifying true brain interaction from EEG data using the imaginary part of coherency. Clinical neurophysiology: official journal of the International Federation of Clinical Neurophysiology 115:2292–2307.

22. Oldfield RC (1971) The assessment and analysis of handedness: the Edinburgh inventory. Neuropsychologia 9:97–113.

23. Oostenveld R, Fries P, Maris E, Schoffelen JM (2011) FieldTrip: Open source software for advanced analysis of MEG, EEG, and invasive electrophysiological data. Computational intelligence and neuroscience 2011:156869.

24. Pain MT, Hibbs A (2007) Sprint starts and the minimum auditory reaction time. J Sports Sci 25:79–86.

25. Polich J (2007) Updating P300: an integrative theory of P3a and P3b. Clin Neurophysiol 118:2128–2148.

26. Popov T, Kastner S, Jensen O (2017) FEF-controlled alpha delay activity precedes stimulus-induced gamma-band activity in visual cortex. The Journal of neuroscience: the official journal of the Society for Neuroscience 37:4117–4127.

27. Proskovec AL, Heinrichs-Graham E, Wiesman AI, McDermott TJ, Wilson TW (2018) Oscillatory dynamics in the dorsal and ventral attention networks during the reorienting of attention. Hum Brain Mapp 39:2177–2190.

28. Rossi S, Huang S, Furtak SC, Belliveau JW, Ahveninen J (2014) Functional connectivity of dorsal and ventral frontoparietal seed regions during auditory orienting. Brain research 1583:159–168.

29. Santangelo V, Olivetti Belardinelli M, Spence C, Macaluso E (2009) Interactions between voluntary and stimulus-driven spatial attention mechanisms across sensory modalities. J Cogn Neurosci 21:2384–2397.

30. Sauseng P, Feldheim JF, Freunberger R, Hummel FC (2011) Right prefrontal TMS disrupts interregional anticipatory EEG alpha activity during shifting of visuospatial attention. Frontiers in psychology 2:241.

31. Sauseng P, Klimesch W, Stadler W, Schabus M, Doppelmayr M, Hanslmayr S, Gruber WR, Birbaumer N (2005) A shift of visual spatial attention is selectively associated with human EEG alpha activity. Eur J Neurosci 22:2917–2926.

32. Shulman GL, Astafiev SV, McAvoy MP, d’Avossa G, Corbetta M (2007) Right TPJ deactivation during visual search: functional significance and support for a filter hypothesis. Cereb Cortex 17:2625–2633.

33. Shulman GL, McAvoy MP, Cowan MC, Astafiev SV, Tansy AP, d’Avossa G, Corbetta M (2003) Quantitative analysis of attention and detection signals during visual search. J Neurophysiol 90:3384–3397.

34. Siegel M, Donner TH, Oostenveld R, Fries P, Engel AK (2008) Neuronal synchronization along the dorsal visual pathway reflects the focus of spatial attention. Neuron 60:709–719.

35. Solis-Vivanco R, Jensen O, Bonnefond M (2018a) Top-down control of alpha phase adjustment in anticipation of temporally predictable visual stimuli. J Cogn Neurosci 30:1157–1169.

36. Solis-Vivanco R, Rodriguez-Violante M, Cervantes-Arriaga A, Justo-Guillen E, Ricardo-Garcell J (2018b) Brain oscillations reveal impaired novelty detection from early stages of Parkinson’s disease. NeuroImage Clinical 18:923–931.

37. Todd JJ, Fougnie D, Marois R (2005) Visual short-term memory load suppresses temporo-parietal junction activity and induces inattentional blindness. Psychol Sci 16:965–972.

38. Tzagarakis C, West S, Pellizzer G (2015) Brain oscillatory activity during motor preparation: effect of directional uncertainty on beta, but not alpha, frequency band. Frontiers in neuroscience 9:246.

39. van Kerkoerle T, Self MW, Dagnino B, Gariel-Mathis MA, Poort J, van der Togt C, Roelfsema PR (2014) Alpha and gamma oscillations characterize feedback and feedforward processing in monkey visual cortex. Proc Natl Acad Sci U S A 111:14332–14341.

40. Van Veen BD, van Drongelen W, Yuchtman M, Suzuki A (1997) Localization of brain electrical activity via linearly constrained minimum variance spatial filtering. IEEE transactions on bio-medical engineering 44:867–880.

41. Vossel S, Thiel CM, Fink GR (2006) Cue validity modulates the neural correlates of covert endogenous orienting of attention in parietal and frontal cortex. NeuroImage 32:1257–1264.

42. Vossel S, Geng JJ, Fink GR (2014) Dorsal and ventral attention systems: distinct neural circuits but collaborative roles. Neuroscientist 20:150–159.

43. WMA (2013) World Medical Association Declaration of Helsinki: ethical principles for medical research involving human subjects. Jama 310:2191–2194.

44. Worden MS, Foxe JJ, Wang N, Simpson GV (2000) Anticipatory biasing of visuospatial attention indexed by retinotopically specific alpha-band electroencephalography increases over occipital cortex. The Journal of neuroscience: the official journal of the Society for Neuroscience 20:RC63.

