## Supplementary figures and images for "New insights on the ventral attention network: Active suppression and involuntary recruitment during a bimodal task"

### Supplementary figure 1

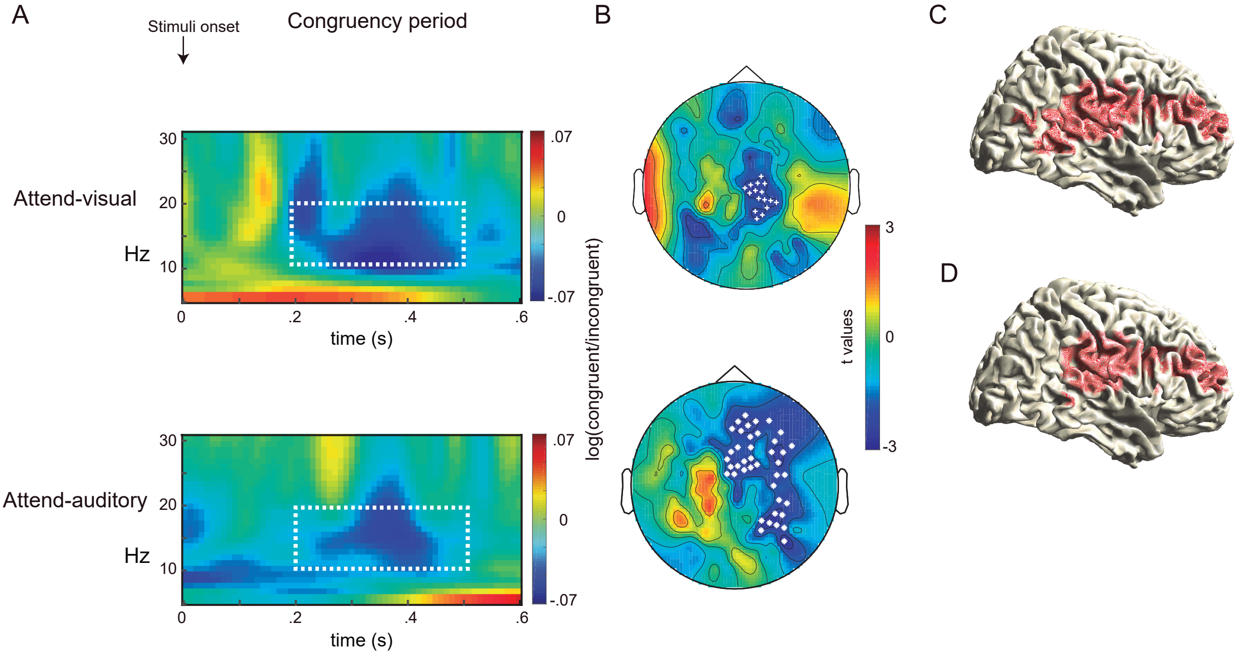

### Supplementary figure 2

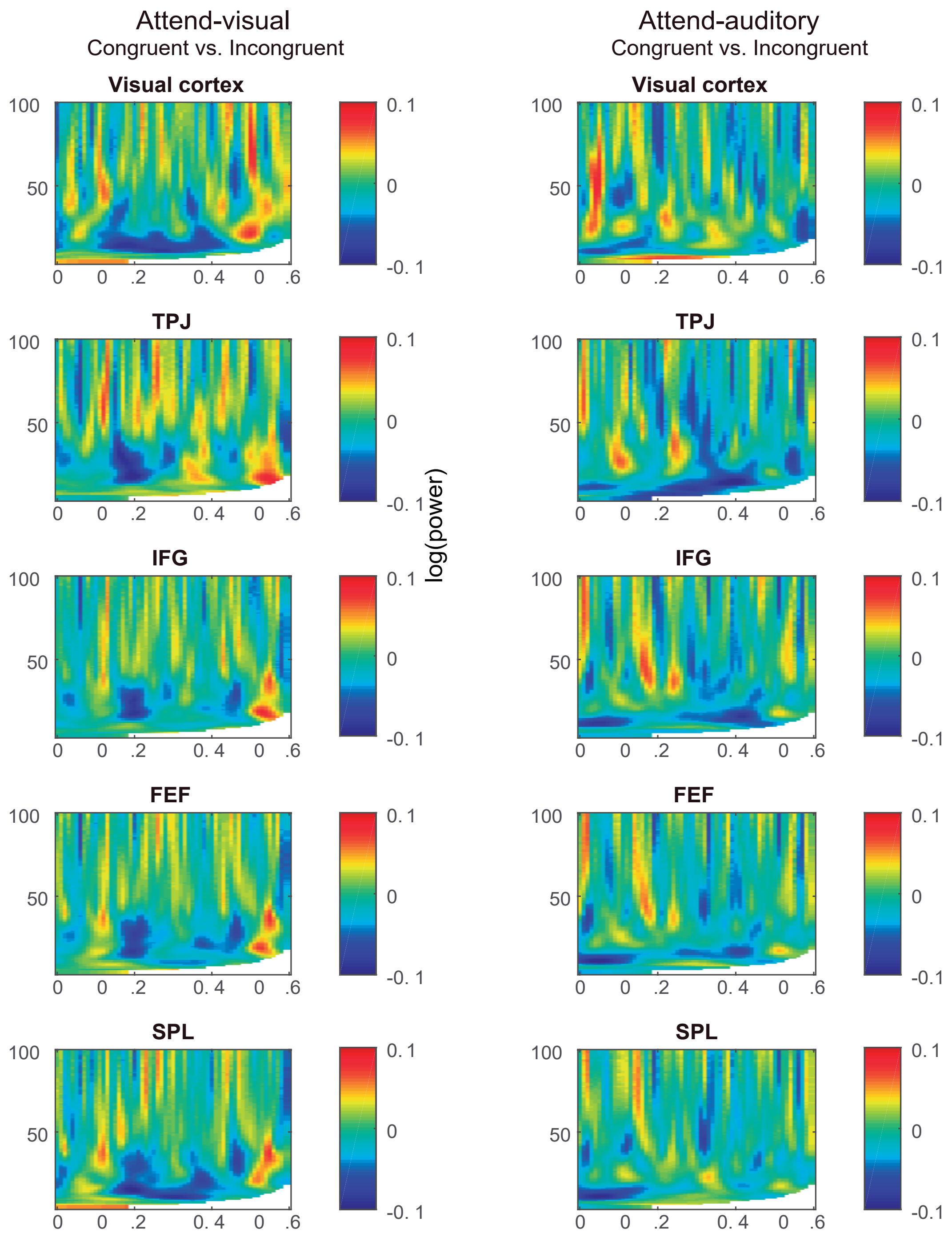
